# Proteasome-dependent degradation of histone H1 subtypes is mediated by its C-terminal domain

**DOI:** 10.1101/2023.06.17.545431

**Authors:** D García-Gomis, J López, A Calderón, M Andrés, I Ponte, A Roque

**Author notes:** Corresponding author: Alicia Roque, Biochemistry and Molecular Biology Department, Biosciences Faculty, Universitat Autònoma de Barcelona, Spain.

## Abstract

Histone H1 is involved in chromatin compaction and dynamics. In human cells, the H1 complement is formed by different amounts of somatic H1 subtypes, H1.0-H1.5 and H1X. The amount of each variant depends on the cell type, the cell cycle phase, and the time of development and can be altered in disease. However, the mechanisms regulating H1 protein levels have not been described. We have analyzed the contribution of the proteasome to the degradation of H1 subtypes in human cells using two different inhibitors: MG132 and bortezomib. H1 subtypes accumulate upon treatment with both drugs, indicating that the proteasome is involved in the regulation of H1 protein levels.

Proteasome inhibition caused a global increase in cytoplasmatic H1, with slight changes in the composition of H1 bound to chromatin and chromatin accessibility and no alterations in the nucleosome repeat length. The analysis of the proteasome degradation pathway showed that H1 degradation is ubiquitin-independent, whereas the whole protein and its C-terminal domain can be degraded directly by the 20S proteasome. Our study shows that histone H1 protein levels are under tight regulation preventing its accumulation in the nucleus. We revealed a new regulatory mechanism for histone H1 degradation, where the C-terminal disordered domain is responsible for its targeting and degradation by the 20S proteasome.

**Statement:** Histone H1 subtypes are a family of proteins involved in the regulation of chromatin structure. This work describes the degradation mechanism controlling the levels of histone H1 subtypes and the region within these proteins involved in the initial recognition. This regulatory mechanism protects the cell nucleus from the damaging effects of its accumulation.

## Introduction

Histone H1 is a multigene family associated with the regulation of chromatin structure. In humans, the H1 family is composed of eleven subtypes or variants. Seven subtypes (H1.0-H1.5, H1X) are differentially expressed in somatic cells, while the remaining four are germ-line specific ^1^. Somatic subtypes are subdivided into two groups: replication-dependent (RD) and replication-independent (RI), according to their expression patterns during cell cycle ^2,3^.

Histone H1 subtypes are basic proteins with three structural domains: the N-terminal domain (NTD), the globular domain (GD), and the C-terminal domain (CTD). The GD has approximately 80 residues and a stably folded ^4,5^. It is responsible for H1 binding to the nucleosome dyad and it is highly conserved in evolution ^4,6^. The NTD is a short domain of 20-36 residues, while the CTD is the longest domain with about 100 residues. Both terminal domains are intrinsically disordered enriched in proline, serine, alanine, and especially lysine^7–11^. The CTD is the main determinant of chromatin compaction within histone H1 ^12^. This domain contains several cyclin-dependent kinases (CDK) consensus sites, which are modified in a cell cycle-dependent manner and affect the secondary structure of the CTD bound to DNA and chromatin, as well as its interaction with chromatin ^13–15^. H1 phosphorylation also facilitates nuclear export ^16^.

The protein levels of histone H1 are tightly regulated during development as they can alter chromatin compaction and transcription. In stem cells, there is approximately one H1 molecule in every two nucleosomes (H1: nucleosome ratio of 0.5), favoring chromatin accessibility and high transcriptional activity ^17,18^. In adults, the H1: nucleosome ratio increases, with values between 0.8 and 1 ^19^. In cells with very low transcriptional activity, like chicken erythrocytes, this ratio increases up to 1.3, promoting gene silencing^20–22^. Gene knockout or knockdown affecting one or two H1 subtypes is not lethal, as the loss is compensated by other subtypes, usually H1.0 ^23,24^. However, triple knockout of H1.2, H1.3, and H1.4 in mice is deleterious during gestation, and the embryonic stem cells derived from these embryos had alterations in the nucleosome spacing and impaired differentiation ^18,23^. Therefore, it is interesting to study the regulatory mechanisms controlling H1 protein levels.

Proteolysis is a crucial regulatory mechanism in the maintenance of proteostasis. Three mechanisms contribute to protein degradation: proteases, the lysosomal system, and the proteasome complex. Proteasomal degradation is responsible for the elimination of most damaged, misfolded, or unfolded proteins in the cell in the nucleus and the cytoplasm ^25^. The proteolytic activity is within a barrel-shaped protein complex known as the 20S proteasome or core particle, which can associate with different regulatory particles on one or both ends ^26^.

There are two pathways of proteasomal degradation of proteins: ubiquitin (Ub)-dependent and Ub-independent. Ub-dependent proteolysis is mediated by the 26S proteasome, formed by the 20S proteasome and the 19S regulatory particle, and the targeting of substrates involves the addition of one or more ubiquitin monomers ^27^. Proteins degraded by the Ub-independent proteasomal pathway are recognized by different signals, such as specific amino acid sequences ^28^, post-translational modifications ^29^, or disordered regions enriched in basic and flexible amino acids ^30^. Degradation is carried out directly by the 20S proteasome or coupled with PA28 or PA200 regulatory particles^31,32^.

Core histones are long-lived proteins, but their degradation is triggered under specific conditions, such as response to EGF, DNA damage, and oxidative stress (reviewed in ^33,34^). In contrast, the proteolytic mechanisms involved in the regulation of histone H1 protein levels are largely unexplored. Early studies showed that the turnover rate of histone H1 is higher than that of core histones, and its magnitude is variable depending on the replicative state of the cells. H1 half-live in rat brain changed from 13 to 112 days between neonate and adult animals, while its value decreased to hours in K562 cells in culture^35,36^. The degradation rate of histone H1 was enhanced after exposure to oxidative conditions by the action of the 20S proteasome. In these conditions, activation of PARP1 accelerated H1 degradation ^36^. Induction of glycoxidation promoted H1 degradation by the nuclear proteasome ^37^. Histone H1 proteolysis is also enhanced by gamma-irradiation and TNF-induced apoptosis ^38,39^.

In the present work, we have studied the contribution of the proteasome to the regulation of histone H1 protein levels. Using proteasome inhibitors, we have analyzed the accumulation of H1 subtypes, the changes in subcellular distribution, and the effect on chromatin. We have also examined the pathway of proteasomal degradation and the contribution of histone H1 structural domains.

## Results

### Effect of proteasome inhibitors in the protein levels of H1 subtypes

We analyzed the role of the proteasome in the control of the protein levels of histone H1 subtypes by inhibition experiments. The proteasome inhibition was assessed using the peptide Suc-LLVY-AMC, which releases the fluorophore after digestion by the chymotrypsin-like activity of the proteasome ^40^. The results showed that treatment with MG132 inhibited proteasome activity at the selected dose (Figure S1).

The effect on the protein levels was analyzed by Western blot. Treatment with MG132 caused an accumulation of β-catenin, used as a positive control ^41^. We observed an accumulation of all the expressed H1 subtypes in T47D and HeLa (Figures 1A and S2). Subtype H1.3 is not detectable in HeLa, while H1.1 cannot be detected in both cell lines. We found a differential accumulation of somatic H1 subtypes with the highest increase in H1.0, followed by H1X (Figure 1B and S2).

**Figure 1:**
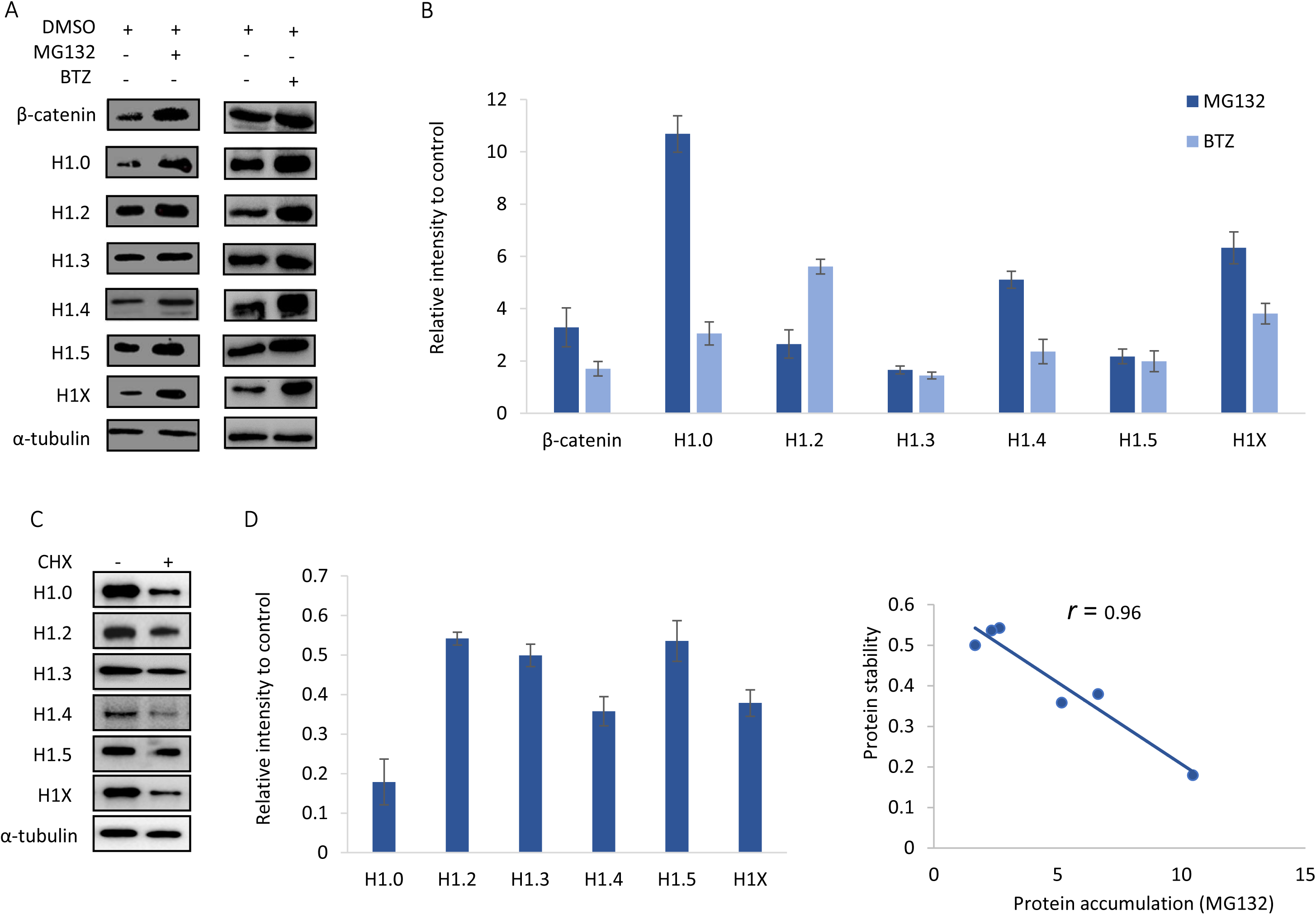
Effects of proteasome inhibition on Histone H1 variants in T47D cells. A. Western Blots of total protein extracts of T47D cells treated with DMSO, MG132 (20μM 12h), and Bortezomib (20nM 12h). C. Western Blots of total protein extract after translation inhibition treatment by cycloheximide (10 μg/ml 8h). B, D. Quantification of the Western blot images of three biological replicates corresponding to A and C, respectively. Error bars correspond to the standard deviation. E. Scatter plot and correlation between protein accumulation and protein stability.

MG132 inhibits the proteolytic activity of the proteasome and calpains, so we also used bortezomib (BTZ), a specific proteasome inhibitor ^26,42^. In the presence of bortezomib, all H1 subtypes accumulated, confirming the role of the proteasome in regulating H1 protein levels (Figure 1A). The level accumulation of H1 subtypes was slightly lower than in MG132, except for H1.2. These results suggested that other proteolytic mechanisms might contribute to the degradation of H1 subtypes.

The distinct accumulation of H1 subtypes upon proteasome inhibition may be explained by differences in protein stability. In T47D, inhibition of protein translation with cycloheximide caused a decrease in the protein levels of H1 subtypes, albeit in different proportions (Figures 1C, 1D). The highest decrease was observed in H1.0, the subtype with the highest accumulation in MG132. Protein accumulation of H1 subtypes upon proteasome inhibition had a negative correlation (r = –0.96) with the protein fraction remaining after translation inhibition (Figure 1E). In HeLa, we observed a similar correlation between the accumulation of H1 subtypes and their stability (r = –0.99) (Figure S3C). In this cell line, H1.0 was also the subtype with the highest accumulation, but its stability could not be measured due to the low levels present at the initial conditions. Our results suggest that H1 subtypes with a higher protein accumulation are less stable, so their protein levels depend on their translation rate.

Protein accumulation may be modulated at the transcript level, so we analyzed the effect of proteasome inhibition in the mRNA levels of H1 subtypes. Treatment with MG132 caused a decrease in the transcript levels of all H1 subtypes in HeLa and T47D (Figure S4). We observed a reduction of more than 40% for the transcripts of H1.0 and H1X and of more than 70% for the replication-dependent subtypes. Considering that H1 transcription is coupled to cell cycle we analyzed if it was affected by treatment with MG132. We found no significant changes in the proportions of cell cycle phases in T47D upon proteasome inhibition (Figure S5). Two main causes could explain the changes described above: i) a feedback regulatory loop triggered by the increase in the protein levels, and ii) indirect effects of the drug.

### Distribution of H1 subtypes after proteasome inhibition

Histone H1 is mainly a nuclear protein, but its presence in the cytoplasm could be detected by immunofluorescence ^43^. The levels detected in the cytoplasm of T47D cells differed depending on the subtype. The estimated percentages of cytoplasmatic H1.0 and H1.5 were less than 10%, approximately 10% for H1.3 and H1.4, and more than 15% for H1.2 and H1X (Figure 2). After treatment with MG132, the amount of all H1 subtypes increased in the cytoplasm, in particular for H1.5 (Figure 2). The increase was statistically significant, except for H1.3. This change in the distribution of H1 subtypes to the cytoplasmatic fraction was confirmed by Western blot (Figure S6). Histone H3 was also detected in the cytoplasm, but at a similar level in both conditions, indicating that the increase in H1 was not due to altered permeabilization of the nuclear membrane or the disruption of chromatin structure (Figure S6). The accumulation in the cytoplasm was observed for all subtypes in T47D cells treated with BTZ (Figure S7) and for H1.2 in HeLa treated with MG132 (Figure S8). The increase was statistically significant, except for H1.0 and H1X in BTZ (Figures S7 and S8).

**Figure 2:**
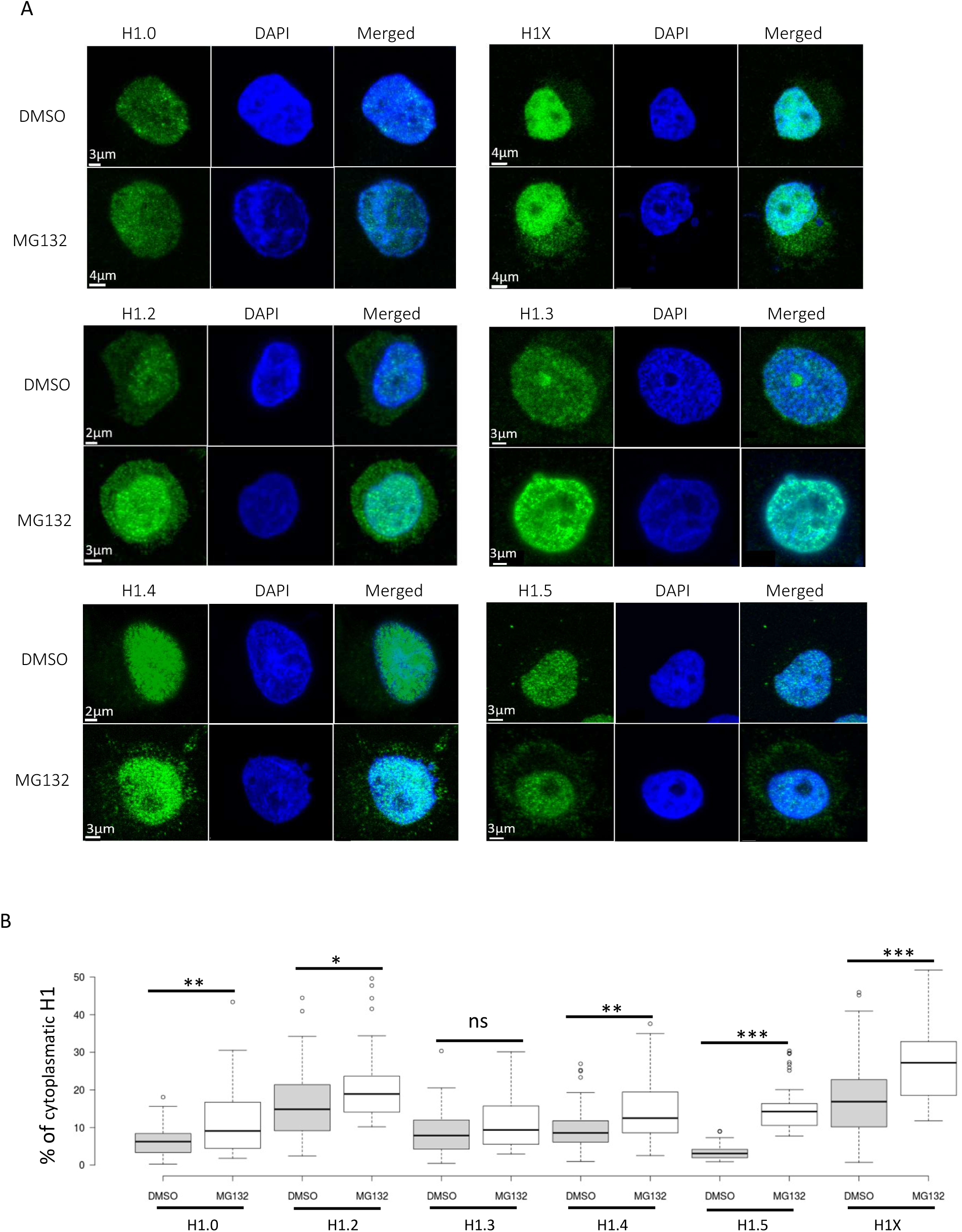
Accumulation of histone H1 subtypes in the cytoplasm of T47D cells after proteasome inhibition with MG132. A. Representative immunohistochemistry images of histone H1 somatic subtypes on T47D cells. The cell nucleus was stained with DAPI. B. Box plots correspond to the quantification of 35-70 cells/subtype and condition. Asterisks denote the p-value of the two-tailed Student’s t-test showing the significance of the difference between untreated and treated cells * p-value < 0.05; ** p-value < 0.01; *** p-value < 0.001; n.s, not significant.

The presence of post-translational modifications (PTMs), in particular phosphorylation, can alter H1 affinity for chromatin, as well as its subcellular localization^16,44^. We used two antibodies against phosphorylated H1 to analyze if there were changes in its subcellular distribution upon proteasome inhibition (Figure 3A). The first antibody recognizes hyperphosphorylated H1, while the second antibody recognizes H1.4T146p. The percentages of phosphorylated H1 in the cytoplasm were higher than 15% and comparable in magnitude to the more abundant subtypes in the cytoplasm. The addition of MG132 caused a significant increase in the cytoplasmic levels of phosphorylated H1 (Figure 3B). The relative increase of both types of phosphorylation suggests that this modification could contribute to the accumulation of H1 in the cytoplasm, although it may not be the only cause.

**Figure 3.**
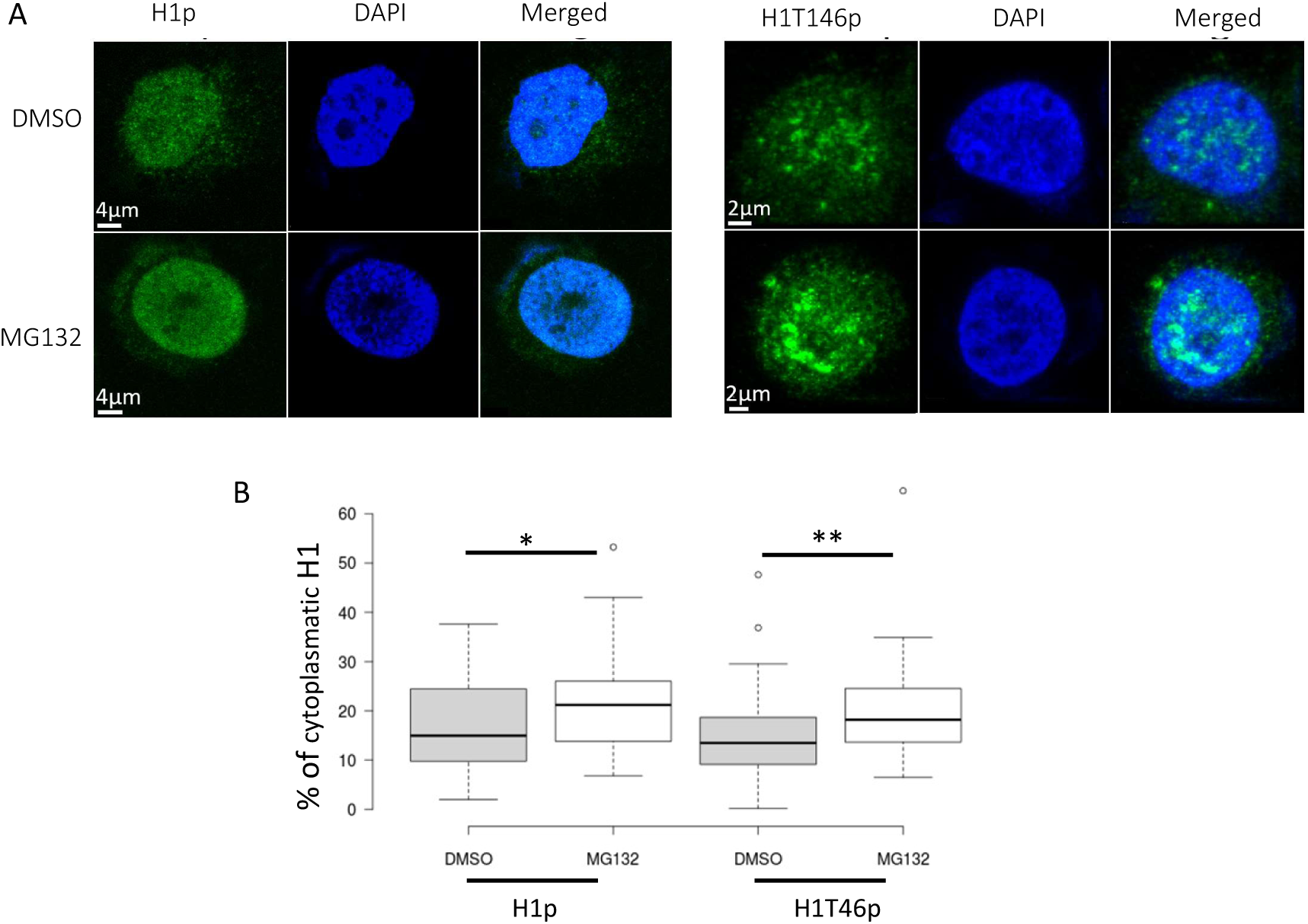
Accumulation of phosphorylated H1 in the cytoplasm of T47D cells after proteasome inhibition with MG132. A. Representative immunohistochemistry images of H1p and H1T56p on T47D cells. Cells were treated with DMSO and MG132 (20 µM 12h). The cell nucleus was stained with DAPI. B. Box plots correspond to the quantification of 35-70 cells/modification and condition. Asterisks denote the p-value of the two-tailed Student’s t-test showing the significance of the difference between untreated and treated cells * p-value < 0.05; ** p-value < 0.01.

### Effects of proteasome inhibition in chromatin

The accumulation of H1 subtypes may alter chromatin structure and compaction. We analyzed the changes in the H1 bound to chromatin after proteasome inhibition using Western blot (Figures 4A and 4B). Treatment with MG132 caused a rearrangement in the proportion of the subtypes bound to chromatin. The amount of H1.0 and H1.2 bound to chromatin increased, while that of H1.4 and H1.5 decreased. Subtypes H1.3 and H1X remained unaltered.

**Figure 4:**
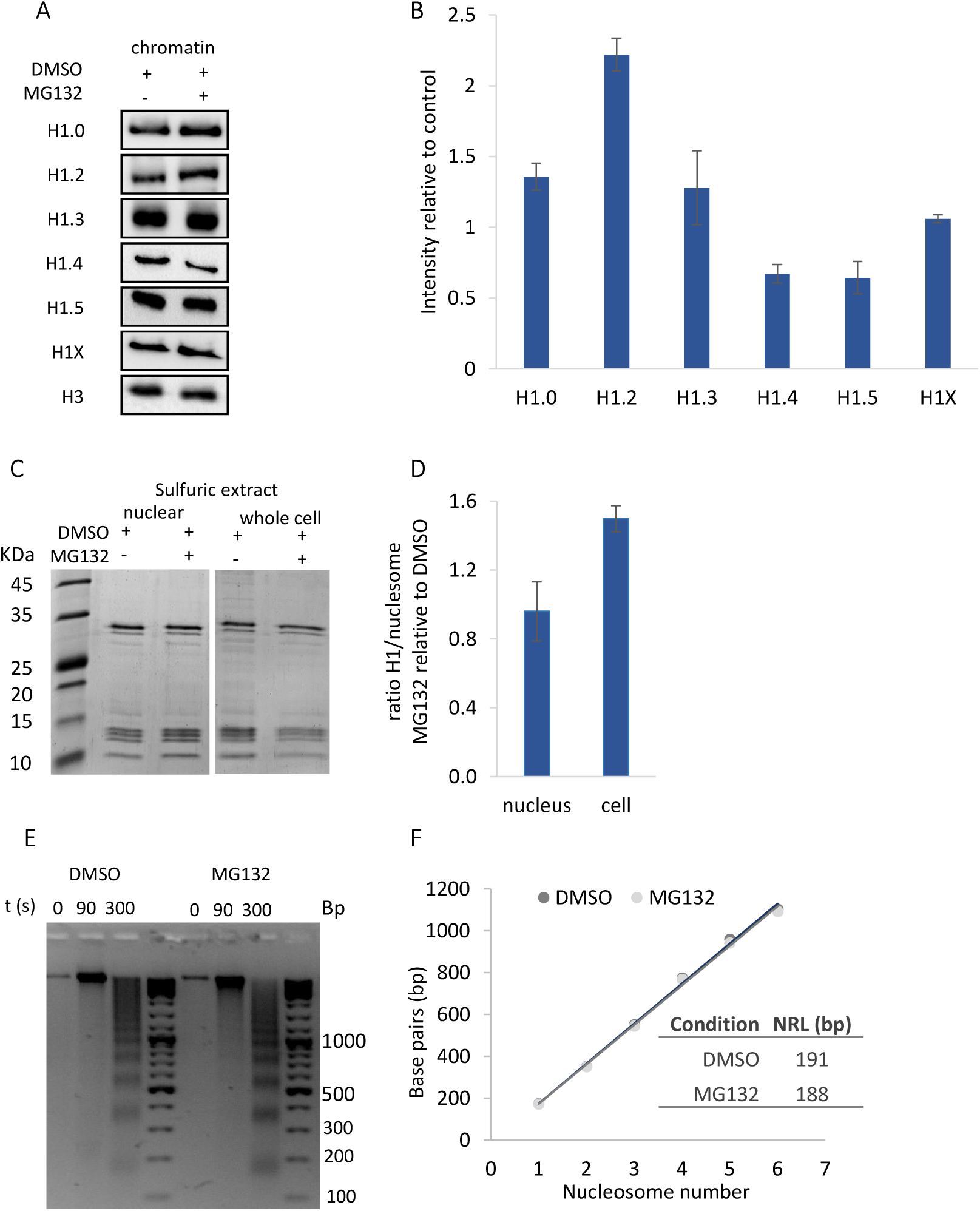
Effects of the accumulation of histone H1 subtypes in chromatin. A. Western Blots and quantification of chromatin-bound H1. B. Quantification of the Western blot images. C. SDS-PAGE of total histone extractions from whole cells and isolated nuclei. D. Fold change of the H1: nucleosome ratio between MG132 and DMSO in whole cells and nuclear extracts. E. Digestion with micrococcal nuclease of chromatin of cells treated with DMSO and MG132. F. Nucleosome repeat length calculated from the digestions shown in E. In all the experiments cells were treated with DMSO as a negative control or with MG132 (20μM 12h). Error bars correspond to standard deviation of three biological replicates.

To assess global changes in the nuclear H1 content we determined the H1: nucleosome ratio using sulfuric acid extractions of the isolated nuclei and whole cells. We found that the ratio H1: nucleosome increased 1.5-fold in the whole cell extract, while in the nucleus it was 0.96-fold, almost identical to the untreated (Figures 4C and 4D). These findings confirmed the increase in the cytoplasm observed by immunofluorescence. We analyzed if there were changes in chromatin accessibility after proteasome inhibition with MG132 by digestion with micrococcal nuclease. The digestion pattern was similar between the two samples, although a slight increase in accessibility could be observed (Figure 4E). However, the nucleosome repeat length (NRL) remained unaltered (Figure 4F). These results suggest that the accumulation of H1 in the cytoplasm after proteasome inhibition prevented significant changes in chromatin structure and compaction.

### Mechanism of degradation of histone H1 subtypes by the proteasome

Protein degradation by the proteasome can occur through different pathways. The most common one is the degradation of ubiquitinated proteins by 26S proteasome. Protein mono– or polyubiquitination causes an increase in the molecular weight of the targeted proteins. In the case of H1, we detected an accumulation of all subtypes after treatment with proteasome inhibitors. However, the electrophoretic mobility of H1s remained unaltered, and no bands of higher molecular weight were observed, suggesting that the contribution of the ubiquitin-dependent proteasome degradation may not be its main degradation pathway.

To prove this hypothesis, we transfected HEK293T cells with a plasmid containing ubiquitin with a histidine tag. The expression of a His-tagged ubiquitin allowed the immunoprecipitation of ubiquitinated proteins, which could be detected by Western blot. First, we confirmed that H1 subtypes were accumulated in HEK293T cells upon proteasome inhibition. In this cell line, the H1 complement, characterized by Western blot, is composed of five subtypes: H1.0, H1.2-H1.4, and H1X. Like in T47D cells, all subtypes increased after treatment with MG132, with H1.0 and H1X being the ones more affected (Figure 5A and 5B).

**Figure 5.**
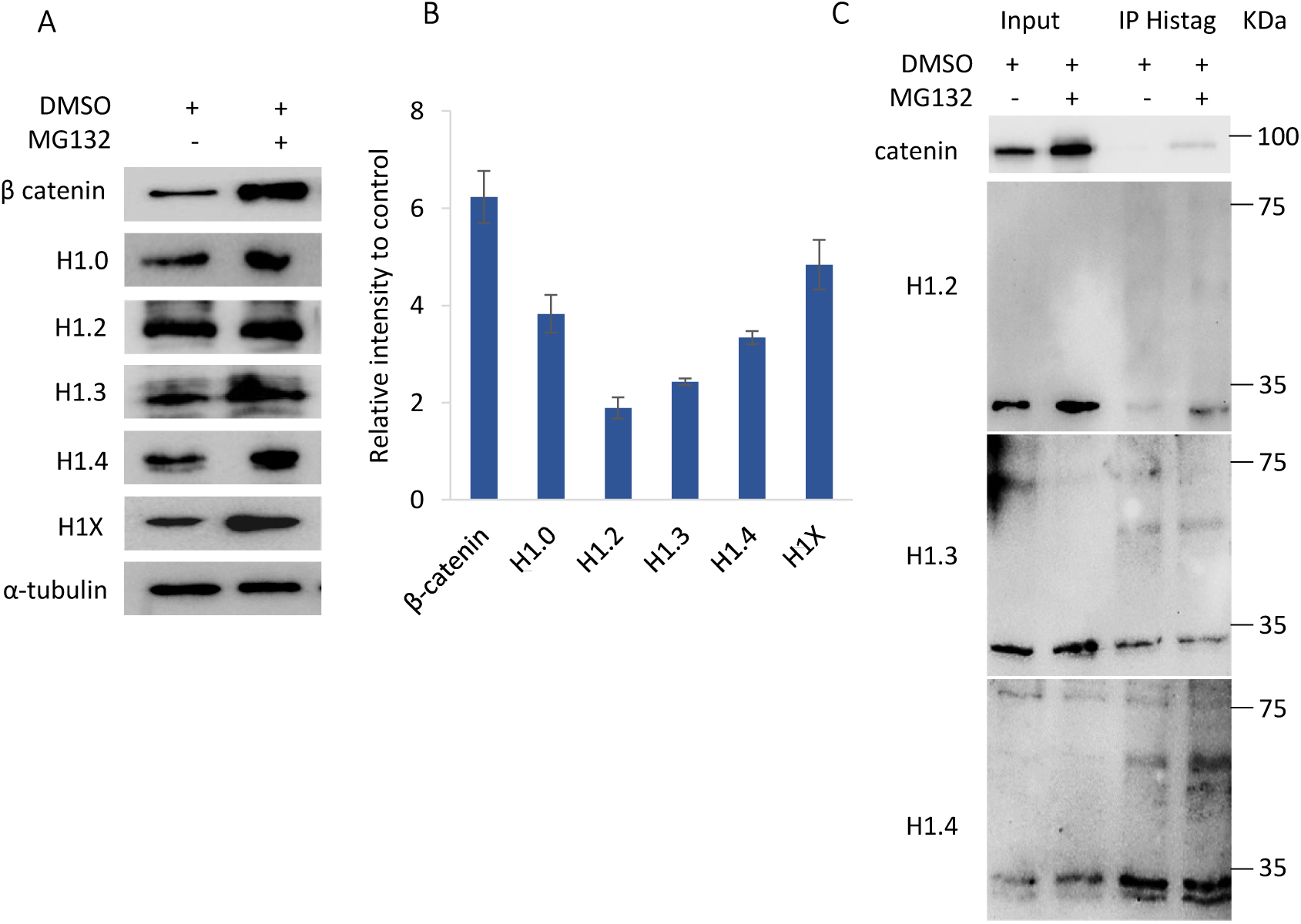
Contribution of the ubiquitin-dependent pathway to H1 degradation. Effects of proteasome inhibition in HEK293T cells. A. Western Blot of total protein extracts upon proteasome inhibition with MG132 (20μM 12h). B. Quantification of the Western blot images of three biological replicates. Error bars correspond to the standard deviation. C. WB of immunoprecipitation of HEK293T cells carrying a tagged ubiquitin and treated with or without MG132.

After immunoprecipitating the protein extract with an antibody against the His-tag, we confirmed by Western blot the presence of ubiquitinated β-catenin in the cells treated with the inhibitor (Figure 5C). We also analyzed the more abundant subtypes expressed in HEK293T. All three subtypes were immunoprecipitated due to unspecific interactions with the His-tag antibody, as the main band detected corresponded to the unmodified protein. We also detected a faint band of higher molecular weight in H1.3 and H1.4. This band was present in the untreated and treated cells showing little or no increase after treatment with MG132 (Figure 5C). However, the increase in the molecular weight of approximately 15-20 kDa without intermediate bands hints that these bands may arise from the unspecific binding of the antibodies or, more likely, from protein aggregates containing H1 subtypes. These results suggest that the 26S proteasome is not the main pathway of H1 subtypes degradation.

Ubiquitin-independent proteasomal degradation can be performed by the 20S proteasome directly or in association with regulatory subunits PA200 or 11S ^26^. The 20S proteasome can directly target basic proteins containing intrinsically disordered domains ^30^. Histone H1 is enriched in basic residues and contains two intrinsically disordered domains, so we analyzed whether the 20S proteasome catalyzed H1 degradation *in vitro*.

Using recombinant H1.0 as a model, we digested the whole protein and its structural domains with the 20S proteasome and analyzed the products by SDS-PAGE. The whole protein was readily digested, with the intact protein disappearing after 60 minutes and detecting lower molecular weight degradation intermediates (Figure 6A). These results suggested that H1 could be degraded directly by the 20S proteasome. The CTD intact band decreased with time, and products of lower molecular weight could be observed, showing similar behavior to the whole protein. In contrast, the GD and bovine serum albumin (BSA) used as a negative control remained stable throughout the reaction (Figure 6A). The NTD was not analyzed because even though it is also intrinsically disordered has only 20 residues, and therefore, it is not suitable for analysis with SDS-PAGE. The whole H1.0, as well as the CTD, is expressed with a histidine tag for purification. To discount the possibility that the tag could be involved in targeting the 20S proteasome, we digested *in vitro* a mixture of native H1s extracted with perchloric acid from T47D. We observed a similar pattern to that of recombinant H1.0, indicating that the degradation of the protein by the 20S proteasome was not associated with the histidine tag.

**Figure 6:**
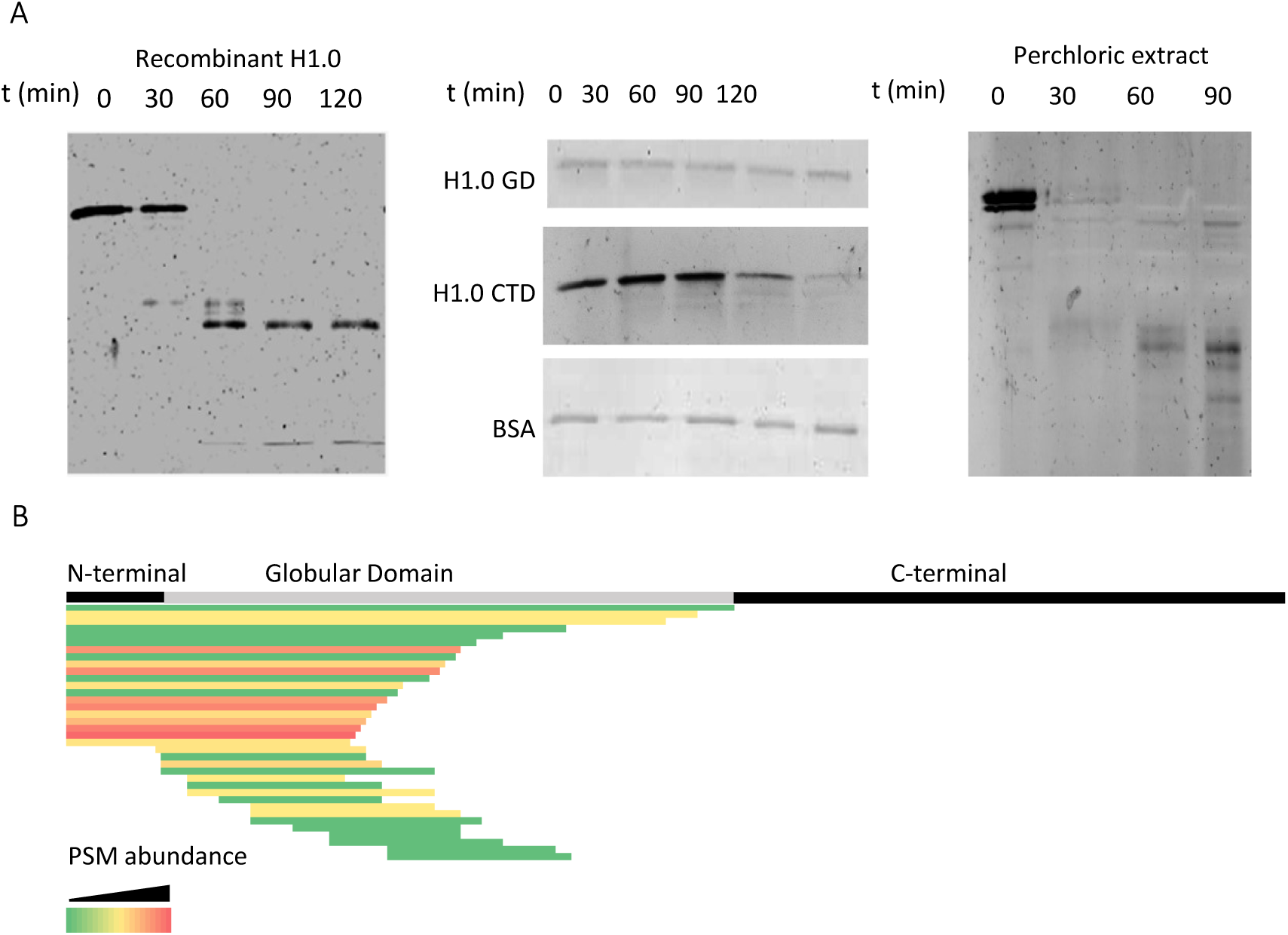
Histone H1 degradation by the 20S proteasome. A. Silver staining of *in vitro* degradation of purified proteins with the 20S proteasome. H1.0, its C-terminal domain (CTD), and its globular domain (GD) are recombinant proteins. Native H1s were obtained by perchloric extraction of T47D cells nuclei. BSA was used as a negative control. B. Schematic representation of the proteoforms of recombinant H1.0 identified by top-down mass spectrometry after partial digestion with the 20S proteasome. Color scale corresponds to the number of PSMs of each proteoform.

To further characterize the contribution of the individual domains of H1.0 to the degradation by the 20S proteasome, we analyzed the digestion intermediates by mass spectrometry (Figure 6B, Table S1). The top-down analysis of partially digested H1.0 allowed the identification of long peptides (> 25 residues), detecting 39 proteoforms of the digested protein with more than one protein-spectrum match (PSM). They could be divided into two groups. The first group had 23 proteoforms, 150 PSMs, and lacked the CTD. The second group had 16 proteoforms with 42 PSMs, lacking the CTD and the NTD. Almost all proteoforms of the second group had two or three PSMs, indicating their low abundance in the sample. These results indicate that the degradation of histone H1 by the 20S proteasome is determined by the intrinsically disordered properties of its CTD.

## Discussion

We have examined the role of the proteasome in the control of histone H1 protein levels. Inhibition of the proteasome with MG132 caused the accumulation of all H1 subtypes in several human cell lines, indicating that it is involved in H1 proteolysis. We observed a negative correlation between H1 accumulation and protein stability. The replication-independent subtypes, especially H1.0, had higher accumulation and were less stable. The regulation of these subtypes at the protein level allows for fast changes in the H1 complement in response to different stimuli. Experimental evidence has confirmed that H1.0 expression is regulated by external stimuli and that this subtype is a key player in the compensation of the H1 levels upon the knockdown of individual or multiple subtypes^3,19,24,45^.

MG132 inhibits the proteasome and cellular proteases like calpains ^26^. For that reason, we analyzed H1 accumulation in the presence of BTZ, a specific inhibitor of the proteasome ^26,42,46^. We found that H1 accumulated less than in MG132, with some changes in specific subtypes. Two explanations, alone or together, could account for our findings: the differences between the two inhibitors and the action of other proteolytic mechanisms. Regarding the first possibility, MG132 at the doses used in our experiments inhibits all three catalytic activities in the proteasome ^47^. Meanwhile, BTZ preferentially inhibits the chymotrypsin-like activity, to a lesser extent, the post-glutamyl activity, and does not inhibit the trypsin-like activity in the standard proteasome ^46^. Considering that histone H1 is enriched in basic amino acids, the trypsin-like activity could be essential for its degradation, explaining some of the differences observed between the inhibitors. As for the second possibility, further experiments will determine whether calpains or other proteases contribute to H1 degradation.

Upon proteasome inhibition with MG132, the transcript levels of H1 subtypes decreased. This difference was not associated with significant changes in the cell cycle. Although the indirect effects of the drug cannot be ruled out, our results open the possibility that transcriptional downregulation could also contribute to preventing the excess of H1 subtypes. The transcriptional regulation of H1 subtypes during the cell cycle and the evidence of co-transcriptional regulation are in favor of this feedback loop ^2,48^. This result suggests that the control of the protein levels of histone H1 is a process with multiple layers.

The multilayered regulation of H1 subtypes at the protein level could be necessary to protect the cell from the cytotoxic effect of its accumulation. An excess of histone H1 can result in its unspecific binding to chromatin, which can alter chromatin structure and transcription. Local alterations in chromatin structure, as well as in the transcript levels were observed in yeast after overexpression of core histones ^49^. In the case of histone H1, some biological systems, such as chicken erythrocytes confirm that a higher content of H1 in the nucleus promotes chromatin compaction and transcriptional silencing ^21,22^. Inducible overexpression of recombinant H1.0 and H1.2 in mouse fibroblasts led to a reduction in the protein levels of the rest of the H1 subtypes while producing a moderate increase in the H1: nucleosome ratio up to 1.3 when H1.0 was overexpressed ^50^. H1 overexpression reduced chromatin nuclease accessibility, increased nucleosome spacing, and caused alterations in gene expression ^51,52^.

Histone H1 is mainly a nuclear protein, but a cytoplasmatic pool of H1 amounting to approximately 10% has been described ^43^. Upon proteasome inhibition, we observed an increase in the cytoplasmatic pool of all H1 subtypes. The accumulation of histone H1 in the cytosol has been observed in mouse models of prion and Alzheimer’s diseases ^44^ and in response to the CDK inhibitor flavopiridol in primary chronic lymphoid leukemia cells ^53^. The preferential accumulation of histone H1 in the cytoplasm, maintaining the H1: nucleosome ratio almost constant in the nucleus, prevented global changes in chromatin structure, as observed in the nuclease accessibility assay. However, local alterations are still possible, as we detected rearrangements in the proportions of H1 subtypes bound to chromatin. Changes in the nuclear chromatin-bound H1 complement and the proportions of H1 subtypes in the cytoplasm could result from the combined effect of their differential characteristics, including protein stability, nuclear localization, chromatin affinity, and PTMs ^3,24,54,55^. In addition, the accumulation of H1 in the cytoplasm could cause unspecific binding to RNA molecules, potentially altering RNA-associated functions. The propensity of H1 to bind RNA molecules and promote phase separation has been recently reported, although its relevance in our conditions was not explored ^56^.

It is known that phosphorylation of H1 promotes its accumulation in the cytoplasm ^16^. We found a slight increase in phosphorylated H1 in the cytoplasm, suggesting that this modification could contribute to its accumulation in this compartment. The extent of the contribution of phosphorylation could be higher than the one we observed, considering that the available antibodies didn’t target directly phosphorylated positions in H1.5, the subtype with the greatest increase in the cytoplasm. Other PTMs that decrease H1 affinity for chromatin like acetylation or parylation, together with phosphorylation at other positions could also play a role in H1 localization to the cytoplasm ^54,55^. However, it is reasonable to think that a part of the H1 in the cytoplasm corresponds to newly synthesized proteins that are not transported to the nucleus.

The ubiquitin-proteasome system controls the metabolism of more than half of the intracellular proteins ^57^. This pathway controls the protein levels of some core histone variants in response to EGF stimulation, SIRT1 overexpression, and during mitosis in embryonic cells ^58–60^. In contrast, the accumulation of H1 subtypes upon proteasome inhibition didn’t result in the appearance of high molecular weight bands corresponding to monoubiquitinated or polyubiquitinated H1s. Immunoprecipitation of His-tagged ubiquitin confirmed that the contribution of this proteolytic mechanism to the control of H1 subtypes is almost nonexistent.

Histone H1 has already been described to be degraded by the 20S proteasome in response to oxidative stress ^36,37^. However, the degron for its recognition by 20S or the proteolytic mechanisms controlling its degradation in normal conditions have not been described. One of the signals for protein targeting to 20S proteasome involves intrinsically disordered regions enriched in basic amino acids ^30^. We found that 20S proteasome can degrade directly recombinant and endogenous H1 subtypes, as well as its CTD. Proteomic analysis of partially digested H1 showed the presence of long peptides lacking the CTD, suggesting that this domain acts as a direct proteasome signal. These results suggest that this mechanism is involved in the targeting and degradation of H1 in normal conditions. Moreover, the increase of the levels of the 20S in certain stress conditions could promote the degradation of damaged H1, among other proteins, preventing chromatin proteotoxicity^61^.

It has been described that intrinsically disordered regions bind to the 20S proteasome, sometimes aided by the presence of basic residues ^30,61,62^. Cryo-EM experiments have shown that the initial binding induced structural alterations consistent with the opening of the central pore in the α-ring and the entering of the substrate to the proteolytic chamber^61^. The opening in the α-ring was wide enough to allow the entry of partially unfolded small globular proteins like ubiquitin and DHFR, when fused to IDR long enough to reach the proteolytic chamber that provided the pulling force for the degradation of the rest of the chimeric protein^30,61^. In the case of H1, its long intrinsically disordered CTD may serve as an initiation site for the recognition by the 20S proteasome with its evenly distributed lysine residues playing an important role in the initial interaction^30^. Binding of the CTD would induce the opening of the gate of the 20S proteasome, allowing the entry and degradation of the rest of the protein. Our results show that cellular proteins composed of intrinsically disordered and stably folded domains can be completely degraded by the 20S proteasome, highlighting the importance of this proteolytic pathway in the maintenance of proteostasis.

## Conclusions

Histone H1 protein levels are tightly regulated by proteasome degradation and, probably, by transcriptional downregulation. Upon proteasome inhibition, the total amount of H1 within the nucleus remained unchanged, while its excess accumulated in the cytoplasm. The alteration in the subcellular distribution minimized the changes in chromatin structure and transcription that could arise from H1 accumulation within the nucleus. We described, for the first time, that H1 intrinsically disordered CTD acts as a primary determinant for its targeting and degradation by the 20S proteasome.

## Methods

### Cell culture and treatments

All the cell lines were grown at 37°C and 5%CO2 in their specific culture media supplemented with 10% fetal calf serum (Gibco) and 1% penicillin-streptomycin (Ddbiolab). Human embryonic kidney 293T cells (HEK293T) and cervical carcinoma cells (HeLa) were cultured in DMEM Glutamax (Corning). For human breast cancer cells (T47D), the culture media was RPMI (Corning), supplemented with 2 mM L-glutamine (Ddbiolab). After harvesting, cells were counted in an automated cell counter (Bio-Rad). For proteasome inhibition, cells were treated with 20 µM MG132 (Sigma) or 20 nM bortezomib (BTZ) (Labnet) for 12 h. Both drugs were dissolved in dimethyl sulfoxide (DMSO). For translation inhibition, cells were treated with 10 μg/mL cycloheximide (Sigma) for 8 h.

### Analysis of proteasomal activity

Proteasome inhibition was assessed by measuring the chymotrypsin-like activity using the fluorogenic peptide, Suc-LLVY-AMC, as described by ^40^. Fluorescence scans were recorded in a Varian Cary Eclipse fluorimeter using excitation wavelength, 345 nm, and emission wavelength interval, 360-500 nm. As a positive control, Suc-LLVY-AMC was digested *in vitro* with 1 unit of chymotrypsin for 15 minutes at 37°C. The chymotrypsin activity was calculated using the fluorescence values at 438 nm and expressed as a percentage of the Suc-LLVY-AMC digested *in vitro*.

### Preparation of protein extracts

Cells were harvested by trypsinization, washed twice with phosphate buffered saline (PBS). For preparing total protein extracts, cells were resuspended in a 50 mM Tris pH 8 buffer containing 300 mM NaCl, 1 % Triton X-100, 0.5 % sodium deoxycholate, 3mM DTT, and a protease inhibitor cocktail (PIC), and incubated for 30 minutes on ice. The mixture was centrifugated at 16000xg, 30 minutes, at 4 °C. The suspension was passed through a needle until it was homogeneous, and centrifugated at 16000xg, 30 minutes, at 4 °C. The supernatant contained a mixture of cellular proteins.

For obtaining cytoplasmatic and nuclear fractions, cells were resuspended in 1 0mM HEPES pH 7.9, 1.5 mM MgCl2, 10 mM KCl and PMSF 0.1 mM. Cell membrane was lysed in a Dounce homogenizer. The nuclei were pelleted by centrifugation at 800 g for 5 minutes. The supernatant, while containing other H1-free organelles, was enriched in cytoplasmatic proteins.

For obtaining linker or total histones acid extraction was used. Perchloric extraction of linker histones in the nuclear fraction was performed as previously described ^63^. Histone sulfuric extraction of whole cells or the nuclear fraction was performed using 0.2 M sulfuric acid, instead of perchloric acid with the same procedure.

All the protein extracts were analyzed by SDS-PAGE and stored at –20°C. Protein concentration was determined with Bradford.

### Western blot

Equivalent amounts of total proteins (10 μg) were separated in a 12% denaturing polyacrylamide gel electrophoresis (SDS-PAGE) and electrotransferred to a polyvinylidene difluoride membrane (PVDF) (EMD Millipore) at 100V for 1h. Immunoblot analyses were performed with the conditions recommended by the manufacturer for the primary and secondary antibodies (Table S2). Specificity of primary antibodies have been validated using knock-down cell lines^64^. Blots were visualized with Clarity Western ECL substrate (Bio-Rad) in a Chemidoc imaging system (Bio-Rad). Band intensities were quantified using Image Lab software (Bio-Rad). α-tubulin was used as loading control.

### RT-qPCR

Total RNA was purified from one million cells with the High Pure RNA Isolation kit (Roche) following the manufacturer’s instructions. Purified RNA was quantified by Nanodrop (Thermo Scientific), and 100 ng were retrotranscribed with iScriptTM cDNA synthesis kit (Bio-Rad) using random hexamers as primers. H1 subtypes and GAPDH were amplified by qPCR using primers specific (Table S3). Fold change was calculated using the ddCT method^65^.

### Immunofluorescence

Cells were grown on the desired conditions, counted, and diluted in PBS to a final concentration of 5·10^4^ cells/mL. They were spun down for 10 min at 500xg in a Thermo Shandon Cytospin 3 using a single-chamber Cytospin funnel as described by ^24^. Cells were air-dried for 1h at room temperature (RT) and then fixed with 2% paraformaldehyde at RT for 10 min. Fixed cells were permeabilized with PBS-0.5 % Triton X-100 and blocked with PBS 0.1 % Tween, 2% bovine serum albumin for 1h at RT. For immunodetection, primary antibodies were incubated at 4°C overnight and secondary antibodies at RT for 1 h (Table S2). Nuclei were stained with 4’, 6-diamidino-2-fenilindol (DAPI) at 0.1 µg/ml. Images were taken using a Leica SP5 AOBS confocal microscope. Two images per slide of three biological replicates, amounting 35-70 cells per condition, were analyzed with Imaris for Cell Biologist package (Oxford instruments). This application allows fluorescence quantification within predefined 3D surfaces. For each cell the fluorescence intensity of H1 was quantified in the 3D surface of the whole cell and that of the nucleus. The nuclear surface was determined using DAPI as a marker. A relative fluorescence score for cytoplasmatic H1 was calculated using the difference between total and nuclear fluorescence of individual cells, normalizing the acquisition parameters for the measurements to be comparable. All cells were visually evaluated to discard artifacts. The values were expressed as a percentage of the total fluorescence. Kolmogorov-Smirnov test was used to determine the adjustment of the values to a normal distribution. The differences between the samples were assessed with the Student’s T-test. All the statistical analyses were performed at https://www.socscistatistics.com/.

### Chromatin extraction

To study H1 subtypes bound to chromatin in untreated cells and after treatment with MG132, chromatin fragments were prepared by sonication. Cells were fixed using 1% formaldehyde, harvested, and sonicated in a Bioruptor (Diagenode) to generate chromatin fragments between 200 and 500 bp. The cross-linking was reversed by incubation at 65°C overnight, and the proteins were analyzed by Western blot.

### Nuclease accessibility assay

The nuclear pellet, obtained as described above, was washed in a buffer containing 10 mM Tris pH 7.4, 3 mM MgCl_2_, 60 mM KCl, and 1% thiodiglycol, centrifuged at 800xg for 5 minutes, and resuspended in the same buffer to a final DNA concentration of 1 mg/mL. Chromatin digestion was performed by adding 0.5 mM CaCl_2_ and micrococcal nuclease (Sigma) at a concentration of 1U/50 µg of DNA at 37°C. Digestion was stopped adding EDTA up to 10 mM. Nuclei were sedimented at 4500xg for 5 minutes and lysed in Tris-EDTA buffer (TE) pH 8.0 for 30 minutes at 4°C. Soluble chromatin was obtained after centrifugation at 16000xg for 10 minutes. Proteins in the supernatant were digested with 100 µg/mL proteinase K (Sigma) at 37°C overnight. DNA fragments were extracted with phenol-chloroform-isoamyl alcohol (25:24:1) and analyzed by agarose gel electrophoresis.

### **A**nalysis of ubiquitinated proteins

Transfection of HEK293T cells with pHis-Ubiquitin (Addgene, 31815) was performed by the calcium phosphate-DNA precipitation method ^66^. After 48h, cells were treated with MG132, harvested, and total protein extracts were prepared as described above. Ubiquitinated proteins were immunoprecipitated using magnetic Protein A Dynabeads (Thermofisher) loaded with an antibody against His-tagged proteins (Finetest, FNab00008). Immunoprecipitation was carried out overnight, at 4°C, in 20 mM Tris pH 7.5, 150 mM NaCl, 1 mM EDTA, 1 mM EGTA, 1% Triton X-100, and (PIC) (Thermofisher). Beads were washed three times, and proteins were eluted in 2x SDS sample buffer at 95°C for 10 min. The presence of ubiquitinated H1s was analyzed by Western blot, using β-catenin as a positive control.

### Preparation of recombinant proteins

Recombinant proteins corresponding to H1.0, its globular and C-terminal domains were expressed and purified from *Escherichia coli*, as previously described ^67^.

### *In vitro* digestion with the 20S proteasome

We analyzed the degradation kinetics of 200 ng/condition of recombinant H1.0, CH1.0, and GH1.0, and 1 µg/condition of the perchloric acid purified histone mixture from T47D cells, using the protocol described by ^30^. Bovine serum albumin (Sigma) was used as a negative control. Proteolytic digestion was performed in 20 mM Tris pH 7,5, 20 mM NaCl, 1 mM EDTA, and 1 mM DTT, in the presence of 5 nM of 20S proteasome (Boston Biochem, E-360), at 37°C. The reaction was stopped by adding electrophoresis loading buffer and incubating the samples at 95°C for 5 minutes. The digestion products were analyzed by 12-15% SDS-PAGE and stained with Coomassie blue or silver (Silver stain plus kit, Bio-rad). Images were taken using a Chemidoc imaging system (Bio-Rad). Degradation intermediates were characterized by top-down mass spectrometry.

### Mass spectrometry analysis

Degradation intermediates of H1.0 digested with proteasome 20S *in vitro* were analyzed by top-down proteomics at the IRB Barcelona Mass Spectrometry and Proteomics Core Facility. The sample was cleaned-up using C4-tips (polyLC) and separated by nanoLC with an Acquity UPLC M-Class BioResolve mAb Column (Waters). Proteoforms were eluted using a combination of two eluents: A. H_2_O 0.1% formic acid; B. CH_3_CN 0.1% formic acid in the following proportions: 10% to 50% of B in 120 min + 50% to 85 % in 7 min. The chromatography was performed at a flow rate of 300 nL/min at 60°C. The sample was ionized by nESI, using Advion Triversa Nanomate (Advion BioSciences) as a source at 1.7 kV, 0.5 psi, in a positive mode. MS/MS was performed in an Orbitrap Fusion Lumos™ Tribrid (Thermo Scientific) in a data-dependent mode. The ions were fragmented by ETD, with 10% supplemental energy from HCD. Proteoform identification was performed using Top-Down PSCW database creation from XML inside Proteome Discoverer software (v2.5) (Thermo Scientific). BioPharma Finder v4.0 (Thermo Fisher Scientific) was used to extract averaged mass spectra from detected chromatographic peaks (MS1). Deconvolution was done using the auto Xtract algorithm on resolved m/z charged species. The slice window option was set to a target average spectrum width of 0.1 min. The recombinant H1.0 sequence was introduced to find a match between the experimental and theoretical masses.

## Supplementary material description

**Supplementary file 1: Figure S1. Inhibition of chymotrypsin activity after treatment with MG132 and BTZ.** A, Fluorescence spectra of Suc-LLVY-AMC peptide in solution (continuous line) and digested with chymotrypsin *in vitro* (dashed line). B, Percentage of chymotrypsin activity in three biological replicates of T47D cells grown in presence of DMSO, 20 µM MG132 (MG132), and 20 nM Bortezomib (BTZ) for 12h. The results are expressed as a percentage of the fluorescence emitted at 438nm by Suc-LLVY-AMC peptide digested with chymotrypsin *in vitro*. Error bars correspond to the standard deviation.

**Supplementary file 2: Figure S2. Accumulation of histone H1 subtypes upon proteasome inhibition in HeLa.** A, Western blot of H1 subtypes after treatment MG132, as described in materials and methods. B, Quantification of the Western blot results in three biological replicates normalized by tubulin. Error bars correspond to the standard deviation.

**Supplementary file 3: Figure S3. Protein stability of H1 subtypes in HeLa** A, Western blot of H1 subtypes after treatment cycloheximide (CHX), as described in materials and methods. B, Protein fraction in two biological replicates of each protein remaining after 8h of treatment with CHX. Error bars correspond to the standard deviation. C, correlation between the accumulation after MG132 treatment and protein stability. r, correlation coefficient.

**Supplementary file 4: Figure S4. Changes in the transcript levels of H1 subtypes after proteasome inhibition.** Cells were treated with MG132, as described in materials and methods. RT-qPCR results are expressed as fold change of the relative expression to GAPDH of each transcript in MG132 treated cells respect to the control cells grown in media supplemented with 0.2 % DMSO of three biological replicates. A. T47D. B. HeLa. Error bars represent the standard deviation.

**Supplementary file 5: Figure S5. Analysis of cell cycle after proteasome inhibition in T47D.** Flux cytometry profiling of cells stained with propidium iodine. A. control cells grown in 0.2% DMSO. B. cells treated with MG132, as described in material and methods. C. Proportion of cells in each phase in two biological replicates, expressed as percentages. Error bars represent the standard deviation.

**Supplementary file 6: Figure S6. Accumulation of histone H1 subtypes in the cytoplasm after proteasome inhibition.** Western Blots of a cytoplasmatic protein extract of T47D cells after treatment with MG132 (20μM 12h).

**Supplementary file 7: Figure S7. Accumulation of histone H1 subtypes in the cytoplasm of T47D cells after proteasome inhibition with Bortezomib.** A. Representative immunohistochemistry images of H1 somatic subtypes in T47D cells. Cellular nucleus was stained with DAPI. Cells were treated with DMSO and Bortezomib (20nM in DMSO, 12h). B. Box plots correspond to the quantification of 35-70 cells/variants and condition. Asterisks denote the p-value of the two-tailed Student’s t-test showing the significance of the difference between untreated and treated cells * p-value < 0.05; ** p-value < 0.01; *** p-value < 0.001; n.s, not significant.

**Supplementary file 8: Figure S8. Accumulation of histone H1.2 in the cytoplasm of HeLa cells after proteasome inhibition.** A. Representative immunohistochemistry images of H1.2 in HeLa cells. Cells were treated with DMSO, MG132 (20 µM) and Bortezomib (20 nM) 12h. Cellular nucleus was stained with DAPI. B. Box plots correspond to the quantification of 35-70 cells/variants and condition. Asterisks denote the p-value of the two-tailed Student’s t-test showing the significance of the difference between untreated and treated cells * p-value < 0.05; ** p-value < 0.01; *** p-value < 0.001; n.s, not significant.

**Supplementary file 9: Table S1. Proteoforms of H1.0 digested with the 20S proteasome *in vitro***.

**Supplementary file 10: Table S2. Antibodies used for Western blot (WB) and immunofluorescence (IF).**

**Supplementary file 11: Table S3. Sequence of the primers of H1 subtypes and conditions for quantitative PCR.**

## List of abbreviations

(DAPI): 4’, 6-diamidino-2-fenilindol
(BTZ): Bortezomib
(BSA): Bovine serum albumin
(CTD): C-terminal domain
(Cryo-EM): Cryogenic electronic microscopy
(CDK): Cyclin-dependent kinase
(SDS-PAGE): Denaturing polyacrylamide gel electrophoresis
(DMSO): Dimethyl sulfoxide
(DTT): Dithiothreitol
(E. coli): Escherichia coli
(GD): Globular domain
(HEK293T): Human embryonic kidney 293T cells
(IPTG): Isopropyl β-D-1-thiogalactopyranoside
(NTD): N-terminal domain
(NRL): Nucleosome repeat length
(PBS): Phosphate buffered saline
(PVDF): Polyvinylidene difluoride membrane
(PMSF): Phenylmethylsulfonyl fluoride
(PTMs): Post-translational modifications
(PIC): Protease inhibitor cocktail
(PSM): Protein-spectrum match
(RD): Replication-dependent
(RI): Replication-independent
(TCA): Trichloroacetic acid
(TE): Tris-EDTA buffer
(Ub): Ubiquitin

## Acknowledgements

This work was supported by grants from the Ministerio de Economía y Competitividad (MINECO) BFU2017-82805-C2-2-P and PID2020-112783GB-C22 awarded to Alicia Roque. Daniel García-Gomis is supported by a PhD fellowship (PIF), awarded by the Autonomous University of Barcelona.

## Authors’ contributions

Conceived and designed the experiments: IP, AR; Performed the experiments: DGG, JLG, AC, MA, IP; Analyzed the data: DGG, JLG, IP, AR; Wrote the paper: IP and AR; Funding acquisition: AR. All authors read and approved the final manuscript.

## Figure legends

**Figure S1.**
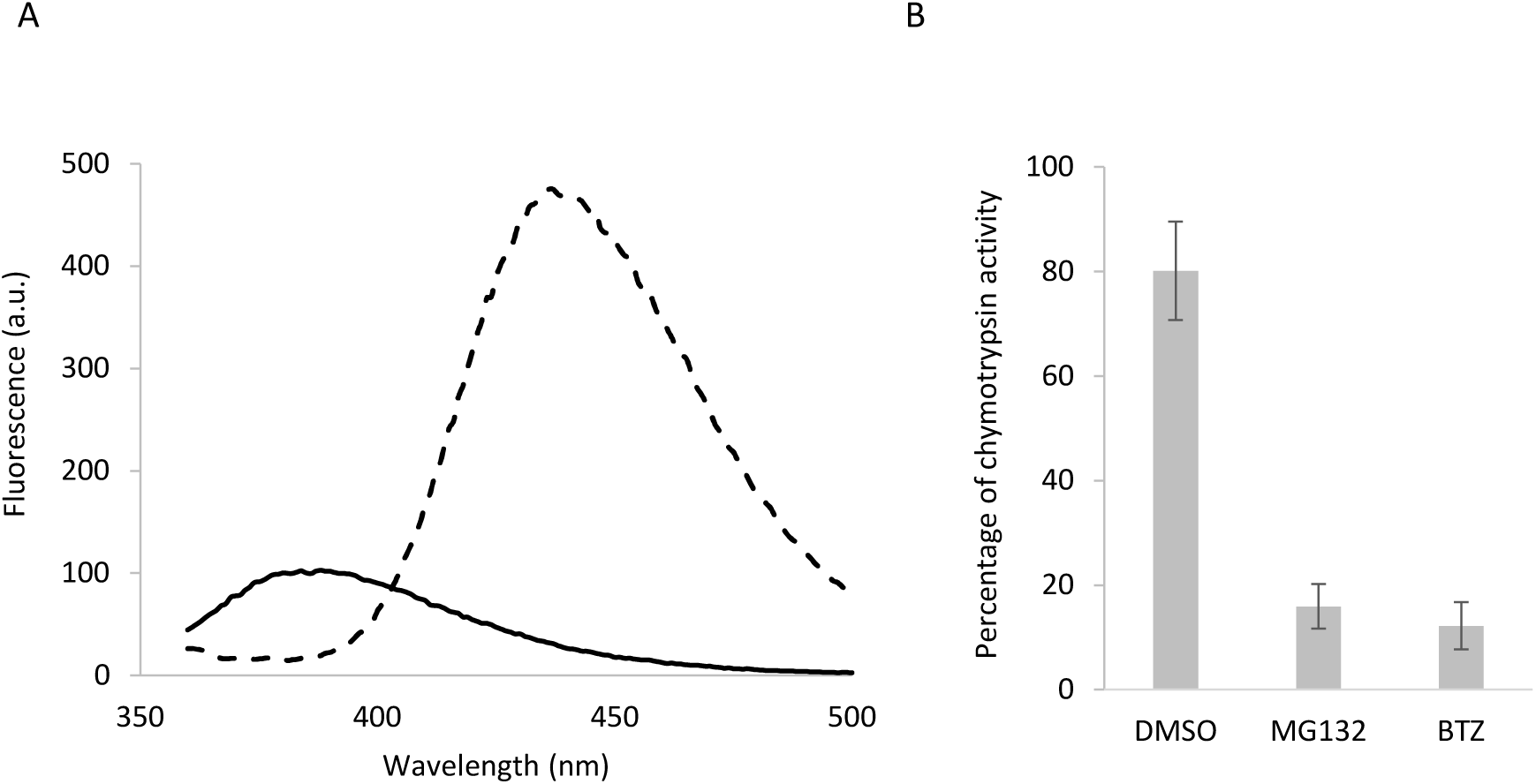
Inhibition of chymotrypsin activity after treatment with MG132 and BTZ. A, Fluorescence spectra of Suc-LLVY-AMC peptide in solution (continuous line) and digested with chymotrypsin in vitro (dashed line). B, Percentage of chymotrypsin activity in three biological replicates of T47D cells grown in presence of DMSO, 20 µM MG132 (MG132), and 20 nM Bortezomib (BTZ) for 12h. The results are expressed as a percentage of the fluorescence emitted at 438nm by Suc-LLVY-AMC peptide digested with chymotrypsin in vitro. Error bars correspond to the standard deviation.

**Figure S2.**
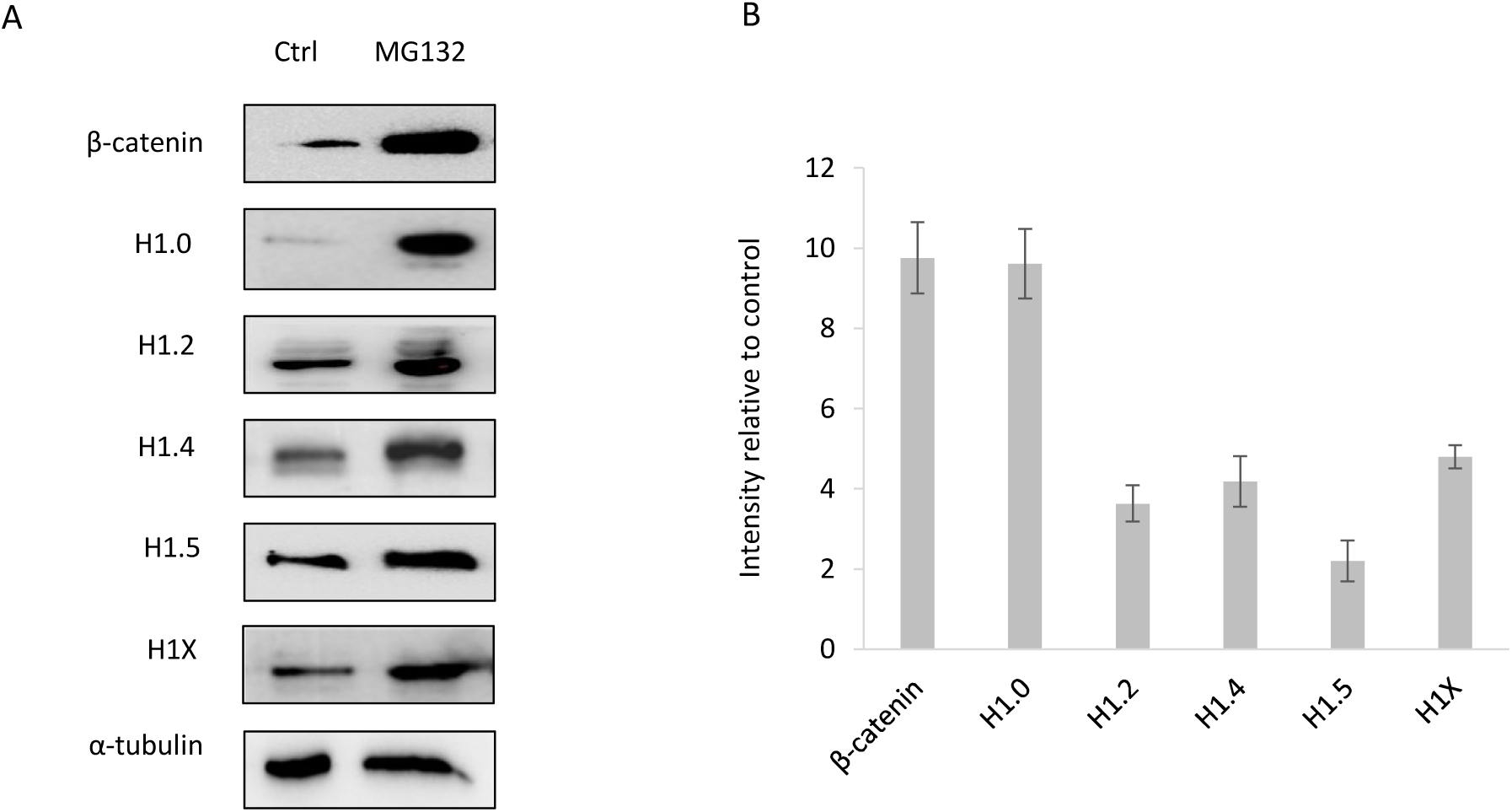
Accumulation of histone H1 subtypes upon proteasome inhibition in HeLa. A, Western blot of H1 subtypes after treatment MG132, as described in materials and methods. B, Quantification of the Western blot results in three biological replicates normalized by tubulin. Error bars correspond to the standard deviation.

**Figure S3.**
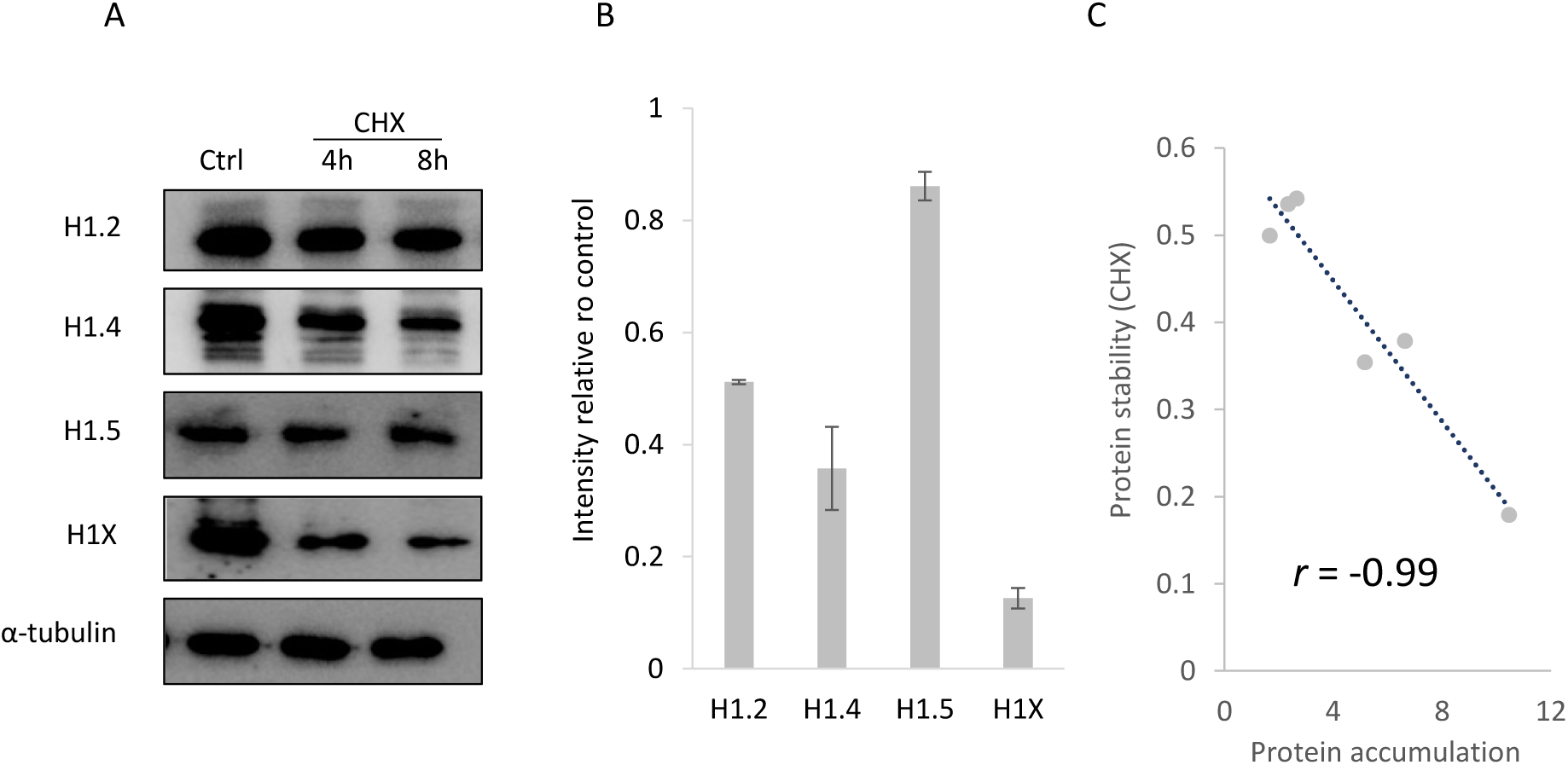
Protein stability of H1 subtypes in HeLa. A, Western blot of H1 subtypes after treatment cycloheximide (CHX), as described in materials and methods. B, Protein fraction in two biological replicates of each protein remaining after 8h of treatment with CHX. Error bars correspond to the standard deviation. C, correlation between the accumulation after MG132 treatment and protein stability. r, correlation coefficient.

**Figure S4.**
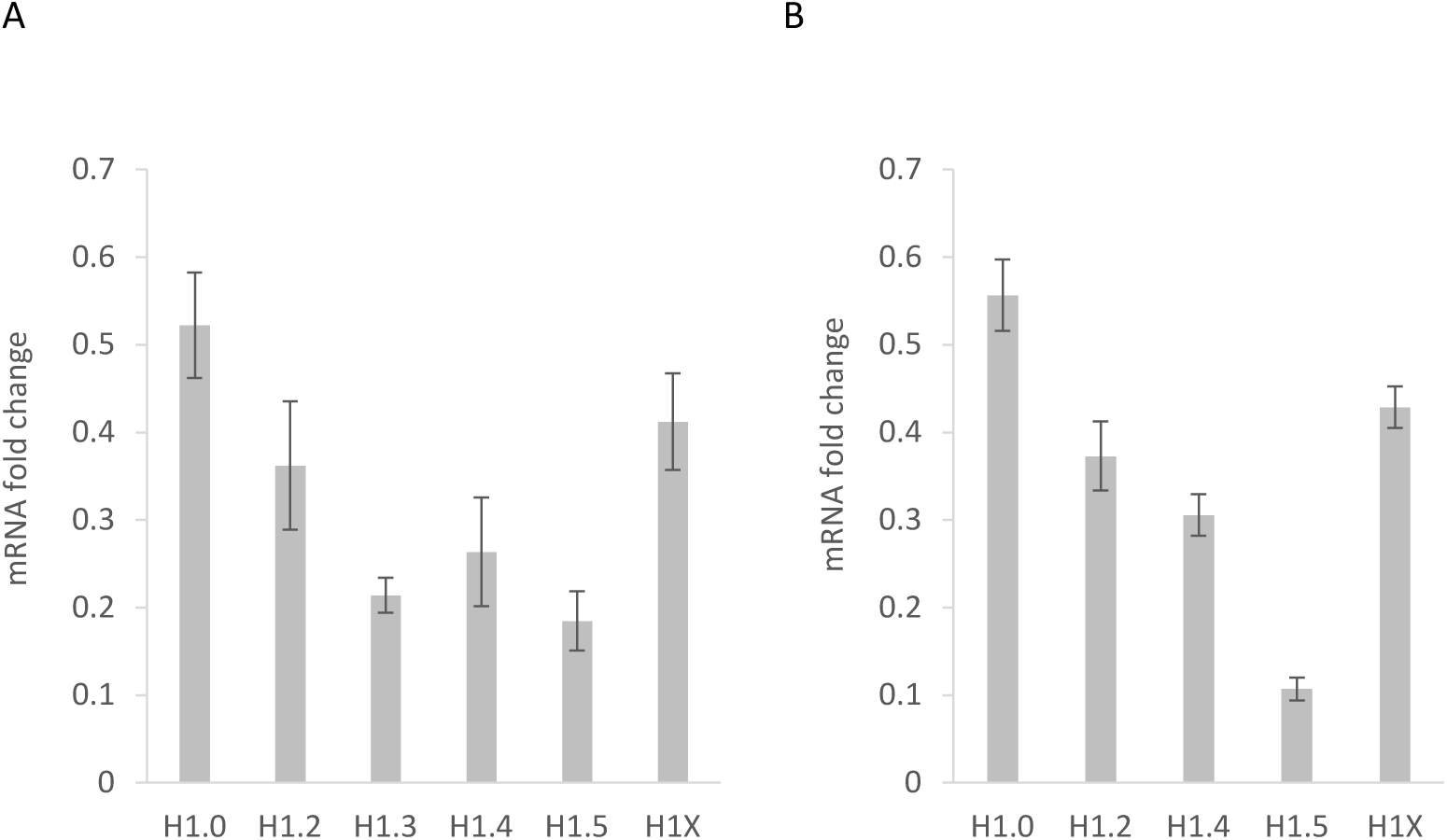
Changes in the transcript levels of H1 subtypes after protesome inhibition. Cells were treated with MG132, as described in materials and methods. RT-qPCR results are expressed as fold change of the relative expression to GAPDH of each transcript in MG132 treated cells respect to the control cells grown in media supplemented with 0.2 % DMSO of three biological replicates. A. T47D. B. HeLa. Error bars represent the standard deviation.

**Figure S5.**
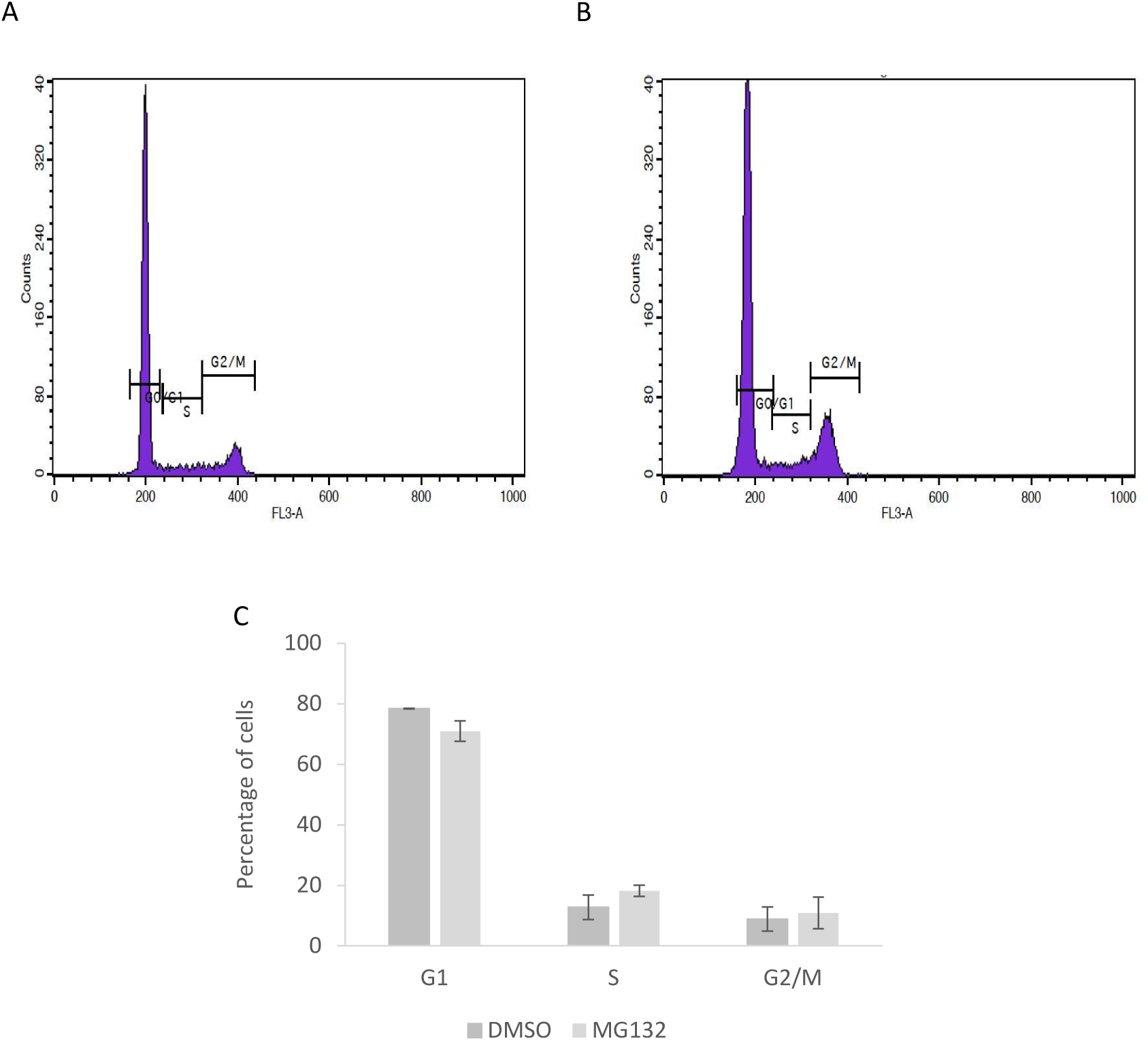
Analysis of cell cycle after proteasome inhibition in T47D. Flux cytometry profiling of cells stained with propidium iodine. A. control cells grown in 0.2% DMSO. B. cells treated with MG132, as described in material and methods. C. Proportion of cells in each phase in two biological replicates, expressed as percentages. Error bars represent the standard deviation.

**Figure S6.**
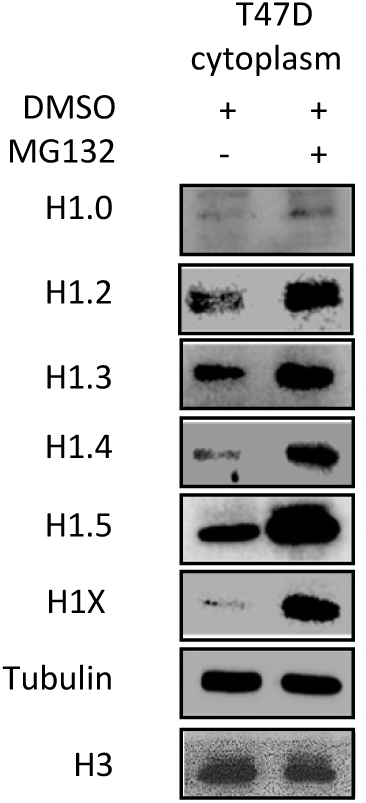
Accumulation of histone H1 subtypes in the cytoplasm after proteasome inhibition. Western Blots of a cytoplasmatic protein extract of T47D cells after treatment with MG132 (20μM 12h).

**Figure S7:**
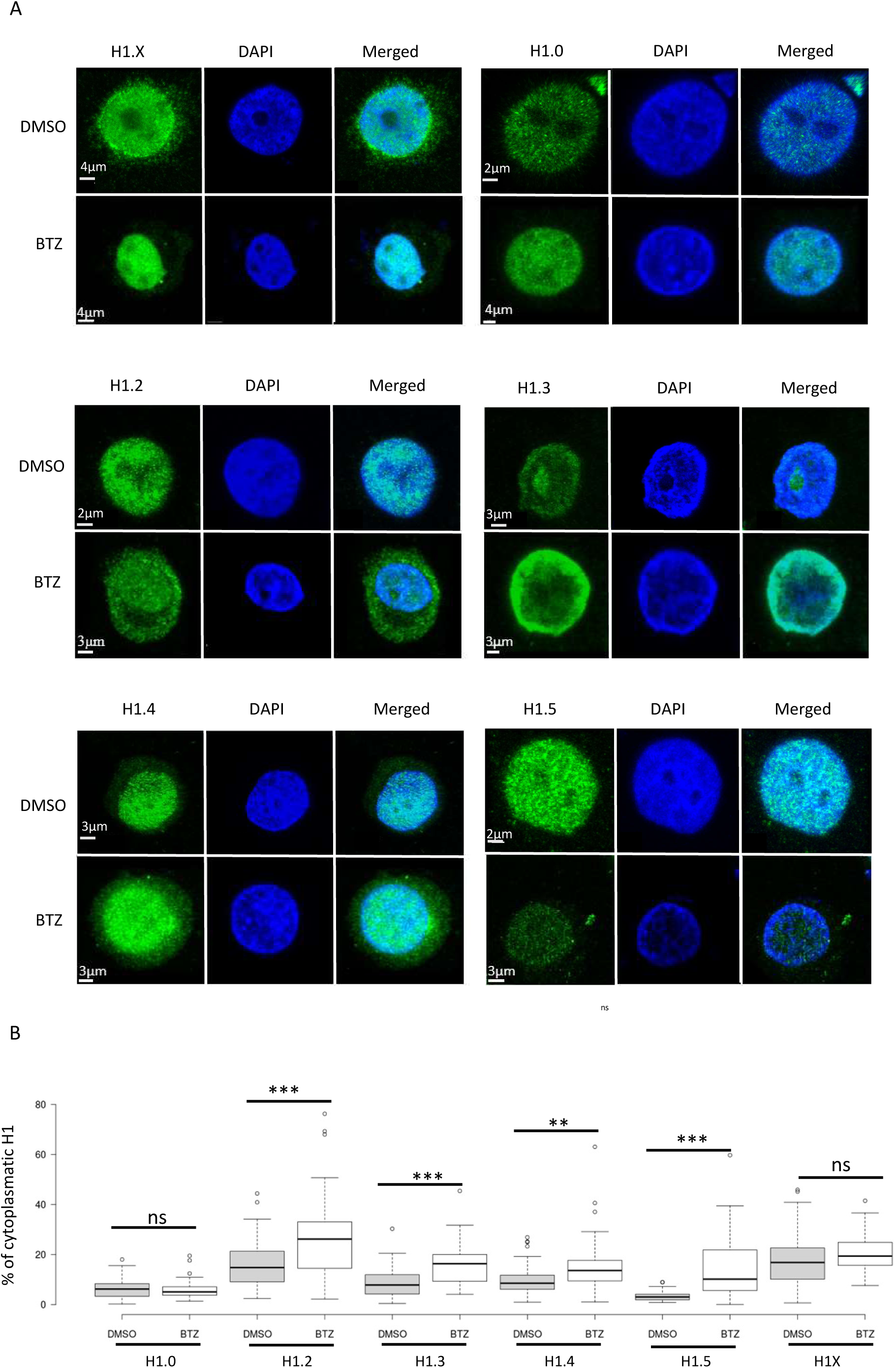
Accumulation of histone H1 subtypes in the cytoplasm of T47D cells after proteasome inhibition with Bortezomib. A. Representative immunohistochemistry images of H1 somatic subtypes in T47D cells. Cellular nucleus was stained with DAPI. Cells were treated with DMSO and Bortezomib (20nM in DMSO, 12h). B. Box plots correspond to the quantification of 35-70 cells/variants and condition. Asterisks denote the p-value of the two-tailed Student’s t-test showing the significance of the difference between untreated and treated cells * p-value < 0.05; ** p-value < 0.01; *** p-value < 0.001; n.s, not significant.

**Figure S8:**
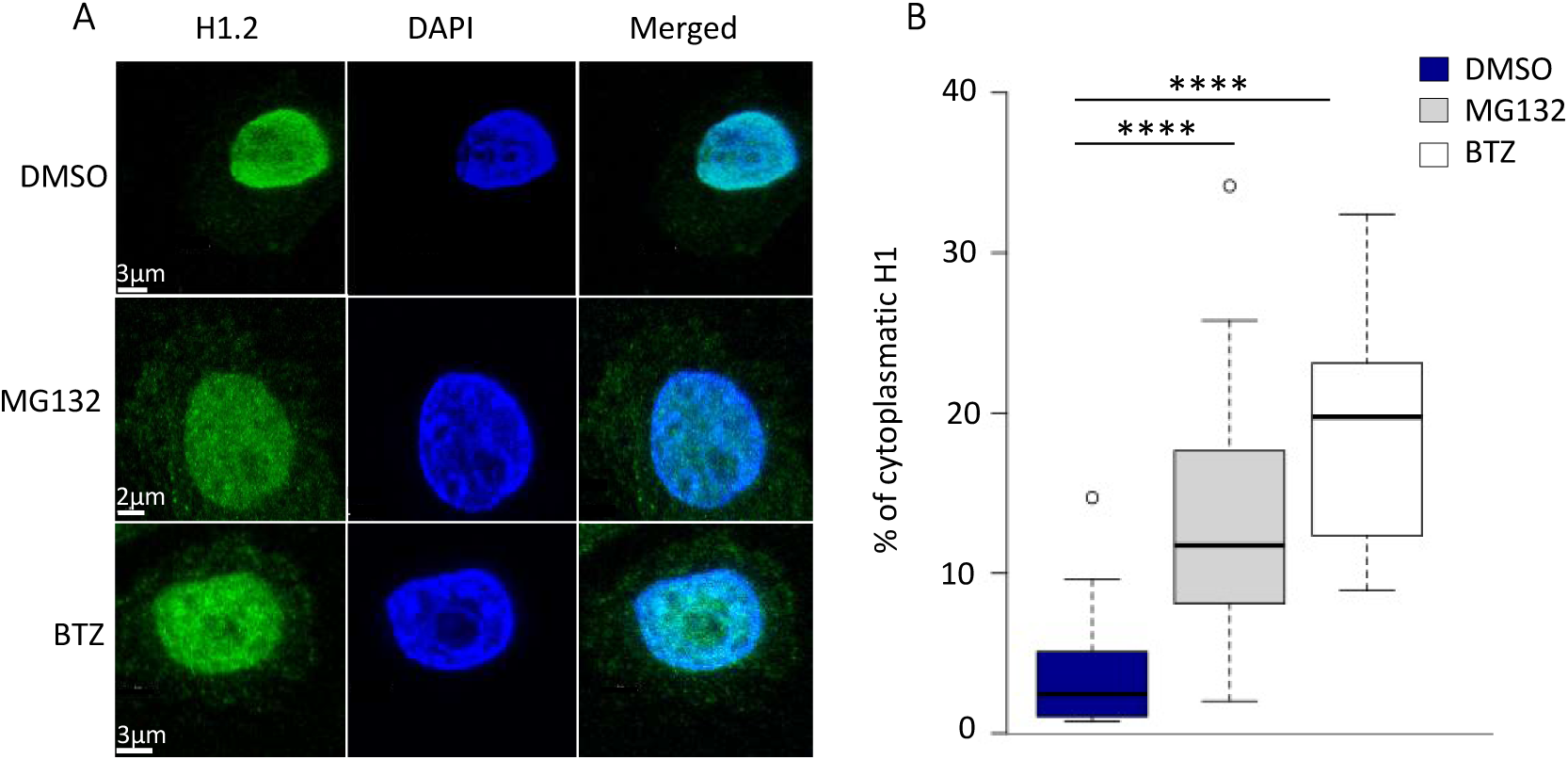
Accumulation of histone H1.2 in the cytoplasm of HeLa cells after proteasome inhibition. A. Representative immunohistochemistry images of H1.2 in HeLa cells. Cells were treated with DMSO, MG132 (20 µM) and Bortezomib (20 nM) 12h. Cellular nucleus was stained with DAPI. B. Box plots correspond to the quantification of 35-70 cells/variants and condition. Asterisks denote the p-value of the two-tailed Student’s t-test showing the significance of the difference between untreated and treated cells * p-value < 0.05; ** p-value < 0.01; *** p-value < 0.001; n.s, not significant.

